# The Cryo-EM Structure of a Pannexin 1 Channel Reveals an Extracellular Gating Mechanism

**DOI:** 10.1101/2019.12.30.890780

**Authors:** Kevin Michalski, Johanna L. Syrjanen, Erik Henze, Julia Kumpf, Hiro Furukawa, Toshimitsu Kawate

## Abstract

Pannexins are large-pore forming channels responsible for ATP release under a variety of physiological and pathological conditions. Although predicted to share similar membrane topology with other large-pore forming proteins such as connexins, innexins, and LRRC8, pannexins have minimal sequence similarity to these protein families. Here, we present the cryo-EM structure of a pannexin 1 (Panx1) channel at 3.0 Å. We find that Panx1 protomers harbor four transmembrane helices similar in arrangement to other large-pore forming proteins but assemble as a heptameric channel with a unique gate formed by Trp74 in the extracellular loop. Mutating Trp74 or the nearby Arg75 disrupt ion selectivity whereas altering residues in the hydrophobic groove formed by the two extracellular loops abrogates channel inhibition by carbenoxolone. Our structural and functional study establishes the extracellular loops as the unique structural determinants for channel gating and inhibition in Panx1 thereby providing the founding model to study pannexins.

## Introduction

Large-pore forming channels play important roles in cell to cell communication by responding to diverse stimuli and releasing signaling molecules like ATP and amino acids (Giaume et al., 2013; Ma et al., 2016; Okada et al., 2018; Osei-Owusu et al., 2018). Pannexins are a family of ubiquitously expressed large-pore forming channels which regulate apoptosis progress (Chekeni et al., 2010), blood pressure (Billaud et al., 2011; Billaud et al., 2015), and neuropathic pain (Bravo et al., 2014; Weaver et al., 2017; Mousseau et al., 2018). While pannexins have limited sequence identity with innexins (∼15% identity), they have virtually no sequence similarity to other large-pore forming channels (Panchin et al., 2000). Among the pannexin family, pannexin 1 (Panx1) has garnered the most attention for its role as a large-pore forming channel responsible for ATP release from a variety of cell types (Bao et al., 2004; Dahl, 2015). Different kinds of stimuli have been reported to activate Panx1 including voltage, membrane stretch, increased intracellular calcium levels, and positive membrane potentials (Bruzzone et al., 2003; Bao et al., 2004; Locovei et al., 2006; Wang et al., 2014; Chiu et al., 2018). Panx1 is also targeted by signaling effectors, such as proteases and kinases, to permanently or temporarily stimulate channel activity (Pelegrin and Surprenant, 2006; Thompson et al., 2008; Sandilos et al., 2012; Billaud et al., 2015; Lohman et al., 2015). The above evidence suggests that Panx1 has a capacity to integrate distinct stimuli into channel activation leading to ATP release. Despite playing critical roles in a variety of biological processes, a mechanistic understanding of pannexin function has been largely limited due to the lack of a high-resolution structure. Here, we show the cryo-EM structure of Panx1, which reveals the pattern of heptameric assembly, pore lining residues, the selectivity filter, and the channel gate.

## Results

### Structure determination and functional characterization

To identify a pannexin channel suitable for structure determination, we screened 34 pannexin orthologues using Fluorescence Size Exclusion Chromatography (FSEC)(Kawate and Gouaux, 2006). Frog Panx1 (frPanx1; 66% identical to human, **Figure1-figure supplement 1**) displayed high expression levels and remained monodisperse when solubilized in detergent, suggesting high biochemical integrity. We further stabilized frPanx1 by truncating the C-terminus by 71 amino acids and by removing 24 amino acids from the intracellular loop between transmembrane helices 2 and 3 (**Figure1-figure supplement 1**). This construct, dubbed “frPanx-ΔLC”, displayed high stability in detergents and could be purified to homogeneity (**Figure1-figure supplement 2a and b**). We verified that frPanx1 forms a functional pannexin channel by whole-cell patch clamp electrophysiology (**Fig.1a and b; Figure1-figure supplement 2e and f**). Purified frPanx1-ΔLC was reconstituted into nanodiscs composed of MSP2N2 (an engineered derivative of apolipoprotein) and soybean polar lipids, and subjected to cryo-electron microscopy (cryo-EM) and single particle analysis (**Figure1-figure supplement 2c and d**). We used a total of 90,185 selected particles for 3D reconstruction at 3.0 Å resolution (**Figure1-figure supplement 3**). The map quality was sufficient for *de novo* model building for the majority of frPanx1-ΔLC with the exception of disordered segments of the N-terminus (residues 1-11), ECL1 (88-100), and ICL1 (157-194) (**Fig. 1c; Figure1-figure supplement 4, and Table 1**).

**Table 1.**
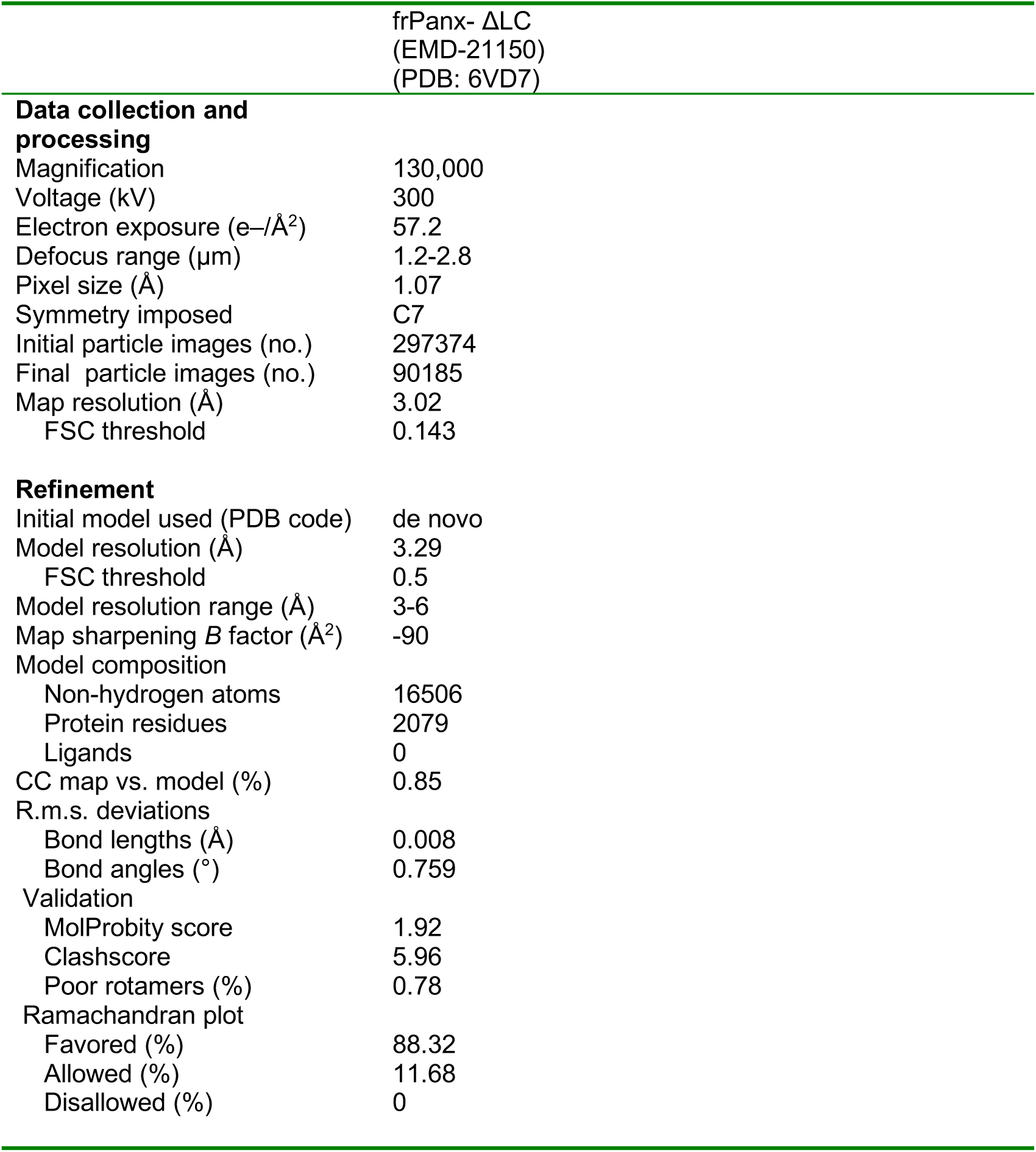
Cryo-EM data collection, refinement and validation statistics.

**Figure 1.**
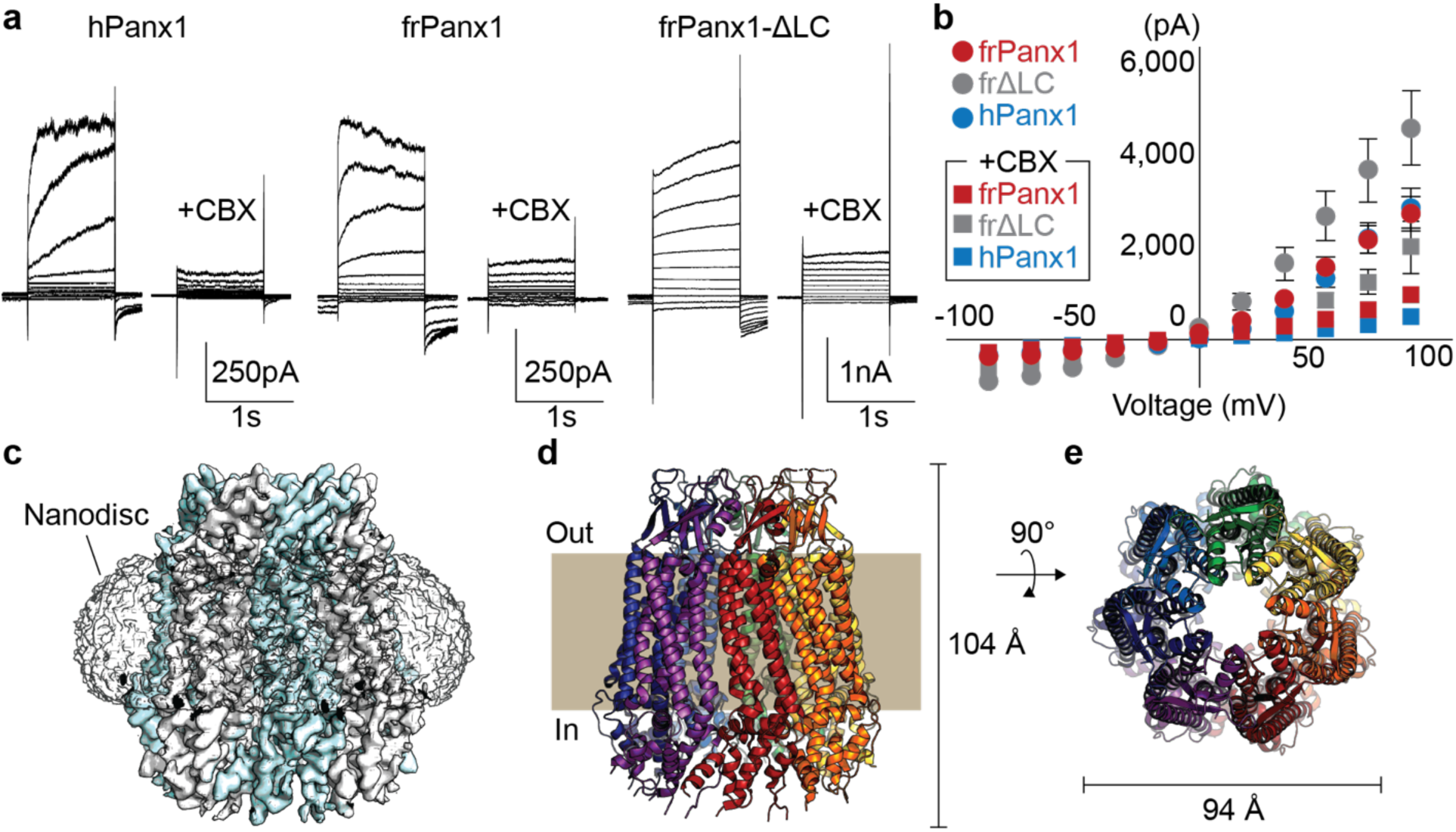
frPanx1 forms a heptameric ion channel. **a**, Whole-cell patch clamp recordings from HEK 293 cells expressing hPanx1, frPanx1, and frPanx1-ΔLC. Cells were clamped at −60 mV and stepped from −100 mV to +100 mV for 1 s in 20 mV increments. To facilitate electrophysiological studies, we inserted a Gly-Ser motif immediately after the start Met to enhance Panx1 channel opening as we have previously described (Michalski et al., 2018). CBX (100 μM) was applied through a rapid solution exchanger. **b**, Current-voltage plot of the same channels shown in **a**. Recordings performed in normal external buffer are shown as circles, and those performed during CBX (100 μM) application are shown as squares. Each point represents the mean of at least 3 different recordings, and error bars represent the SEM. **c**, EM map of frPanx1-ΔLC shown from within the plane of the membrane. Each protomer is colored white or blue. **d**, Overall structure of frPanx1-ΔLC viewed from within the lipid bilayer. Each protomer is colored differently, with the extracellular side designated as “out”. **e**, Structure of frPanx1 viewed from the extracellular face.

### Overall structure and protomer features

The frPanx1-ΔLC structure revealed a heptameric assembly, which is unique among the known eukaryotic channels (**Fig. 1d and e**). Other large-pore forming channels include hexameric connexins (Maeda et al., 2009) and LRRC8s (Deneka et al., 2018; Kasuya et al., 2018; Kefauver et al., 2018), and the octameric innexins (Oshima et al., 2016) (**Figure2-figure supplement 1**). Our result differs from previous studies that suggest hexameric assembly of pannexin based on single channel recordings on concatemeric channel and negative stain electron microscopy (Boassa et al., 2007; Wang et al., 2014; Chiu et al., 2017). The heptameric assembly observed in the current study is unlikely to be caused by the carboxy-terminal truncation because cryo-EM images of the full-length frPanx1 also display clear seven-fold symmetry in the 2D class averages (**Figure2-figure supplement 2a**). Furthermore, 2D class averages of hPanx1 display a heptameric assembly, but not other oligomeric states (**Figure2-figure supplement 2b**). Thus, overall, our data suggests that the major oligomeric state of Panx1 is a heptamer. This unique heptameric assembly is established by inter-subunit interactions at three locations: 1) ECL1s and the loop between β2 and β3; 2) TM1-TM1 and TM2-TM4 interfaces; and 3) α9 helix and the surrounding α3 and α4 helices, and the N-terminal loop from the neighboring subunit (**Figure2-figure supplement 3**). Notably, the majority of residues mediating these interactions are highly conserved (e.g. Phe67 and Tyr111; **Figure1-figure supplement 1**).

The overall protomer structure of Panx1 resembles that of other large-pore forming channels including connexin, innexins, and LRRC8. Like other large-pore forming channels, each Panx1 protomer harbors four transmembrane helices (TM1-4), two extracellular loops (ECL1 and 2), two intracellular loops (ICL1 and 2), and an amino (N)-terminal loop (**Fig. 2a and b**). The transmembrane helices of Panx1 are assembled as a bundle in which the overall helix lengths, angles, and positions strongly resemble the transmembrane arrangements observed in other large-pore channels (**Fig. 2c**). Features in the ECL1 and ECL2 domains appear to be conserved among large-pore channels despite limited sequence similarity (**Fig. 2d-g; Figure2-figure supplement 1**). For example, the Panx1 ECL1 and ECL2 are joined together by two conserved disulfide bonds (Cys66 with Cys267, Cys84 with Cys248) in addition to several β-strands. ECL1 also contains an alpha-helix that extends towards the central pore and forms an extracellular constriction of the permeation pathway. While much of the transmembrane domains and extracellular loops show similarities to other large-pore forming channels, the Panx1 intracellular domains are structurally unique (**Figure2-figure supplement 1**). ICL1 and ICL2, for example, together form a bundle of helices that make contact with the N-terminus. The N-terminus of Panx1 forms a constriction of the permeation pathway and extends towards the intracellular region. The first ∼10 amino acids of the N-terminus are disordered in our structure, but these residues might play a role in ion permeation or ion selectivity (Wang and Dahl, 2010).

**Figure 2.**
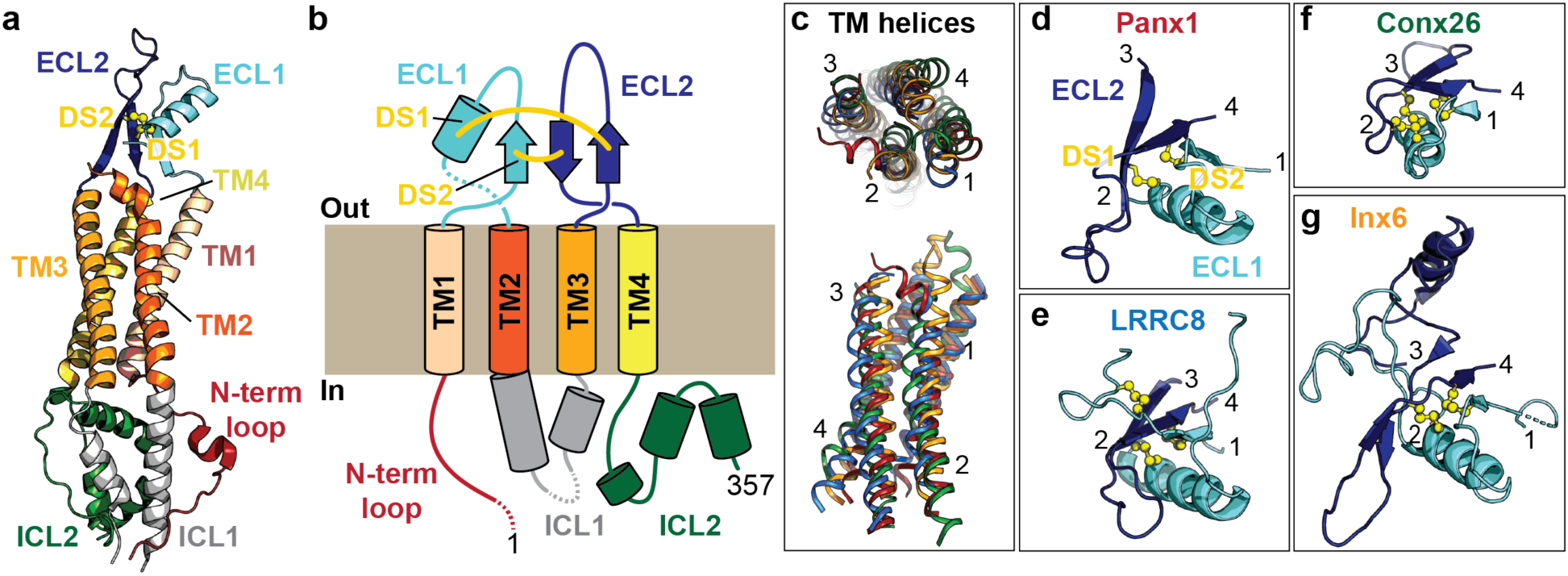
Subunit architecture of frPanx1. **a**, Structure of the frPanx1 protomer. Each domain is colored according to the cartoon scheme presented in **b. c.** Superimposition of the transmembrane helices from frPanx1 (red), connexin-26 (green), innexin-6 (orange), and LRRC8 (blue) shown top-down from the extracellular side (top) or from within the plane of the membrane (bottom). **d-g**, Cartoon representation of the extracellular loops of large pore forming channels. ECL1 is colored in light blue, and ECL2 is colored in dark blue, and disulfide bridges are shown as yellow spheres. These domains are viewed from the same angle (from top) as shown in the top panel in **c**.

### Ion permeation pathway and selectivity

The Panx1 permeation pathway spans a length of 104 Å, with constrictions formed by the N-terminal loop, Ile58, and Trp74 (**Fig. 3a and b**). The narrowest constriction is surrounded by Trp74 located on ECL1, which forms an extracellular gate (**Fig. 3c**). Trp74 is highly conserved among species including hPanx1 (**Figure1-figure supplement 1**). Because Panx1 has been previously characterized as an anion selective channel (Ma et al., 2012; Romanov et al., 2012; Chiu et al., 2014), we wondered if this extracellular gate harbors positively charged amino acids which may contribute to anion selectivity of the channel. Interestingly, Arg75 is situated nearest to the tightest constriction of the permeation pathway (**Fig. 3d**). Though the current structure likely represents a closed conformation based on the lack of channel activity at 0 mV (**Figure2-figure supplement 2e**), we hypothesized that Arg75 might be a major determinant of anion selectivity of Panx1 channels in the open state. To assess whether Arg75 contributes to anion selectivity, we generated a series of point mutations at this position on hPanx1 and compared their reversal potentials (Erev) in asymmetric solutions using whole-cell patch clamp electrophysiology (**Fig. 3e and Figure3-figure supplement 1**). We kept sodium chloride (NaCl) constant in the pipette solution while varying the extracellular solution. When treated with the large anion, gluconate (Gluc^-^), Erev shifted to +26 mV, suggesting the channel is more permeable to Cl^-^ than to Gluc^-^. When exposed to the large cation, *N*-methyl-D-glucamine (NMDG^+^), Erev remained close to 0 mV, suggesting that Na^+^ and NMDG^+^ equally (or do not) permeate Panx1. These results are consistent with Panx1 being an anion-selective channel. The Arg75Lys mutant maintains the positive charge of this position, and displayed Erev values comparable to WT. Removing the positive charge at this position, as shown by the Arg75Ala mutant, diminished Cl^-^ selectivity as the Erev in NaGluc remained near 0 mV. Interestingly, the Erev in NMDGCl shifted to −22 mV, suggesting the channel had lost anion selectivity and Na^+^ became more permeable than NMDG^+^. A charge reversal mutant, Arg75Glu, shifted the Erev in NaGluc to −16 mV and in NMDGCl to −45 mV, indicating that Gluc^-^ became more permeable to Cl^-^. Overall, these results support the idea that the positively charged Arg75 plays a role in anion selectivity of Panx1.

**Figure 3.**
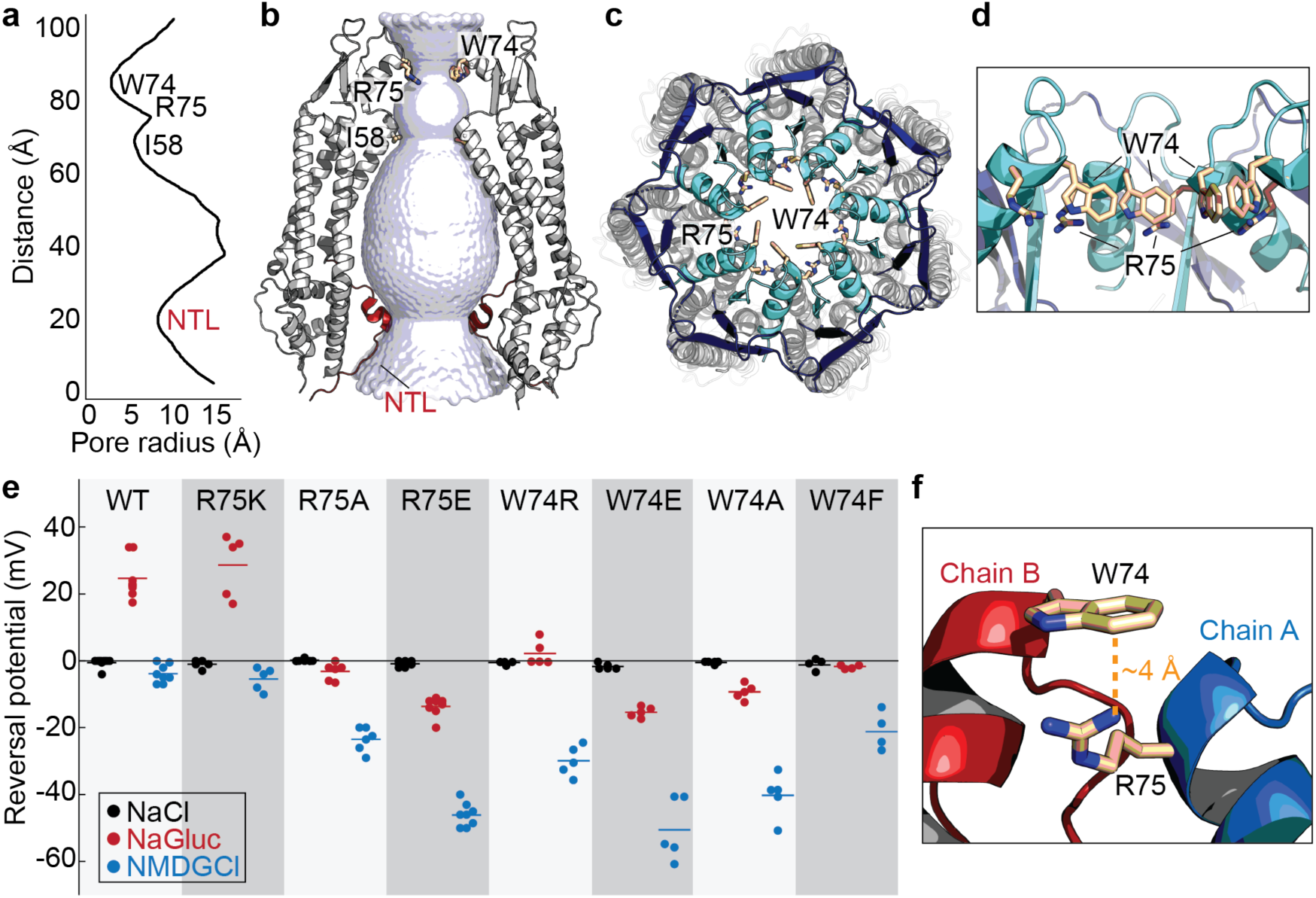
Permeation and ion selectivity of Panx1 channels. **a**, HOLE (Smart et al., 1996) diagram demonstrating constrictions along the permeation pathway. NTL; N-terminal loop. **b**, Surface representation of the internal space along the molecular 7-fold axis running through the center of frPanx1. The surface was generate using HOLE. **c and d**, Top view facing the extracellular side (**c**) or side view (**d**) of frPanx1, with ECL1 shown in light blue and ECL2 in dark blue. Trp74 and Arg75 are shown as sticks. **e**, Reversal potentials of various hPanx1 ion selectivity mutants. Each point represents the Erev measured in NaCl (black), NaGluc (red), or NMDGCl (blue), and bars represent the mean values. I-V curves were obtained by a ramp protocol from −100 mV to +100 mV. **f**, Close-up view of the Trp74-Arg75 interaction at the interface of protomer A (blue) and B (red).

We next wondered if introducing a charge at position 74 might alter ion selectivity of Panx1 channels. Interestingly, both Trp74Arg and Trp74Glu mutants become less selective to anions and more permeable to Na^+^ (**Figure 3e**). These results suggest that introducing a charge at this position disrupts the natural ion selectivity of Panx1 channels but that position 74 itself does not control ion selectivity. We observed that the distance between the guanidyl group of Arg75 and the benzene ring of Trp74 from an adjacent subunit is ∼4 Å, suggesting that these two residues likely participate in an inter-subunit cation-π interaction key to Panx1 ion selectivity (**Figure 3f**). To test this hypothesis, we generated Trp74Ala and Trp74Phe mutations and measured Erev potentials. Trp74Ala showed a marked decrease in Cl^-^ permeability and an increase in Na^+^ permeability, despite preservation of the positive charge at Arg75. A more conservative mutation, Trp74Phe, still disrupted ion selectivity, suggesting that proper positioning of the benzene ring at position 74 is important for anion selection. Altogether, our data suggests that anion selectivity is only achieved when Trp74 and Arg75 form a cation-π interaction. Given that our structure has disordered and truncated regions in the N-terminus, ICL1, and ICL2, it is possible that additional ion selectivity or gating regions exist in the full-length channel. For example, the N-termini of LRRC8 and connexins have been demonstrated to perform an important role in ion selectivity (Kyle et al., 2008; Kronengold et al., 2012; Kefauver et al., 2018). It is possible that the N-terminus of Panx1 is mobile and may further constrict the permeation pathway.

### CBX action mechanism

We have previously demonstrated that CBX, a potent nonselective inhibitor of Panx1, likely acts through a mechanism involving ECL1 (Michalski and Kawate, 2016). In these experiments, mutations at a number of residues in ECL1 rendered Panx1 less sensitive to CBX-mediated channel inhibition. Mapping such residues in the Panx1 structure revealed that they are clustered proximal to the extracellular gate, in a groove formed between ECL1 and ECL2 (**Figure 4a and b**). This supports our previous speculation that CBX is an allosteric inhibitor, not a channel blocker (Michalski and Kawate, 2016).

**Figure 4.**
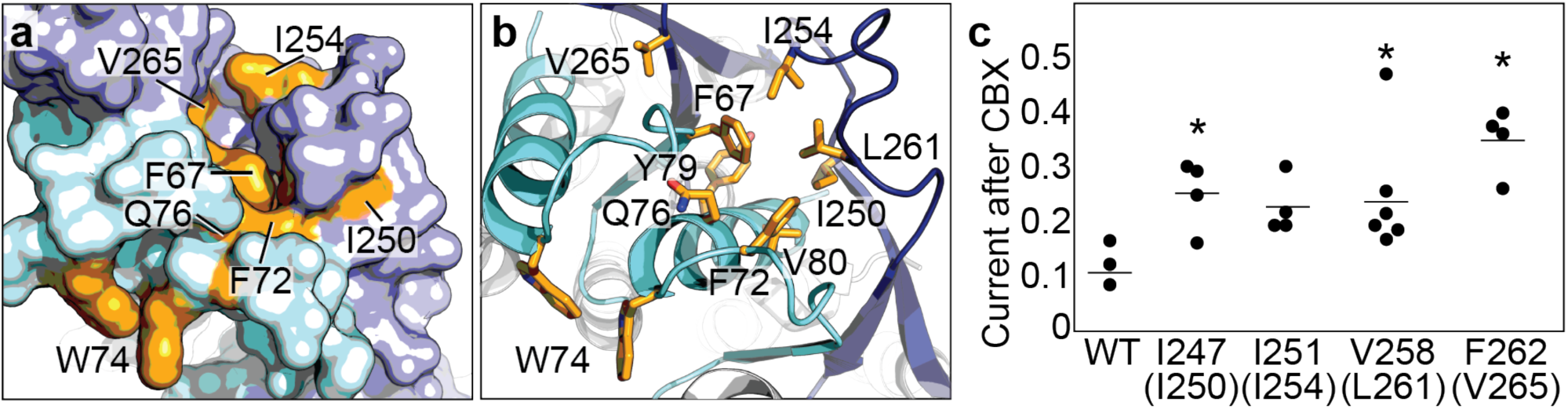
CBX action requires residues from both ECL1 and ECL2. **a and b**, Surface (**a**) and cartoon (**b**) representations of the frPanx1 ECL1 (light blue) and ECL2 (dark blue), with potential CBX-interacting residues shown in orange. **c**, Quantification of whole-cell currents from hPanx1 mutants when treated with CBX (100 μM). Mutants are numbered according to the hPanx1 sequence while the mutants in parenthesis are the corresponding residues in frPanx1. Recordings were performed by stepping to +100 mV in the absence or presence of CBX, and each point represents the normalized current amplitude during the CBX application. Bars represent the mean value from each mutant. Asterisks indicate significance of p<0.05 determined by one-way ANOVA followed by Dunnett’s test comparing WT to each mutant (F262C: p=0.0007; I247C: p=0.0471; V258C: p=0.0363).

Given that this hydrophobic groove is formed also by residues in ECL2, we wondered if residues in ECL2 might also play a role in CBX-mediated inhibition. We mutated selected residues in ECL2 of hPanx1 to cysteines and measured channel activity before and after CBX application. We found that mutations at Ile247, Val258, and Phe262 (hPanx1 numbering) diminished CBX-sensitivity (**Figure 4b**). These data suggest that both ECL1 and ECL2 play important roles in inhibition of Panx1 by CBX. Although we do not have a cryo-EM structure complexed to CBX at this point, we speculate that CBX inhibits Panx channels by binding between ECL1 and ECL2 and ‘locking’ the conformation of gate forming ECL1 in favor of channel closure.

## Discussion

The frPanx1-ΔLC structure uncovered a unique heptameric assembly of a large-pore channel that harbors a novel extracellular gate formed by Trp74 and Arg75. These residues are located on ECL1 and face toward the central pore of the channel and thus, are situated to regulate channel function. The positioning of ECL1 may be regulated by ECL2, which strongly interacts with ECL1 and appears to create a binding pocket for CBX. Both ECL1 and ECL2 may undergo movement based on conformational alterations of the TMDs and cytoplasmic domains. For example, it is conceivable that movement of the TMDs caused by membrane stretch or voltage, or changes in the cytoplasmic domain triggered by caspase cleavage may be coupled to opening and closing of this extracellular gate. The major role of the extracellular gate in function is strongly supported by our experimental results demonstrating that mutating Trp74 and Arg75, as well as surrounding residues in ECL1 and ECL2, alter channel properties like ion selectivity and CBX sensitivity (Michalski and Kawate, 2016). Other domains, especially the N-terminal loop and the C-terminal domain, have been demonstrated to play important roles in Panx1 channel gating (Sandilos et al., 2012; Michalski et al., 2018), suggesting the possibility for another gate on the intracellular side of the channel. While we did not observe a clearly structured gate in other domains, it is possible that they may only be stabilized in particular functional states. Nevertheless, our study provides a fundamental template to conduct further experiments such as cysteine accessibility and molecular dynamics simulations to accelerate our understanding of how this unique large-pore channel functions.

## Figure supplements and Table

**Figure1-figure supplement 1.**
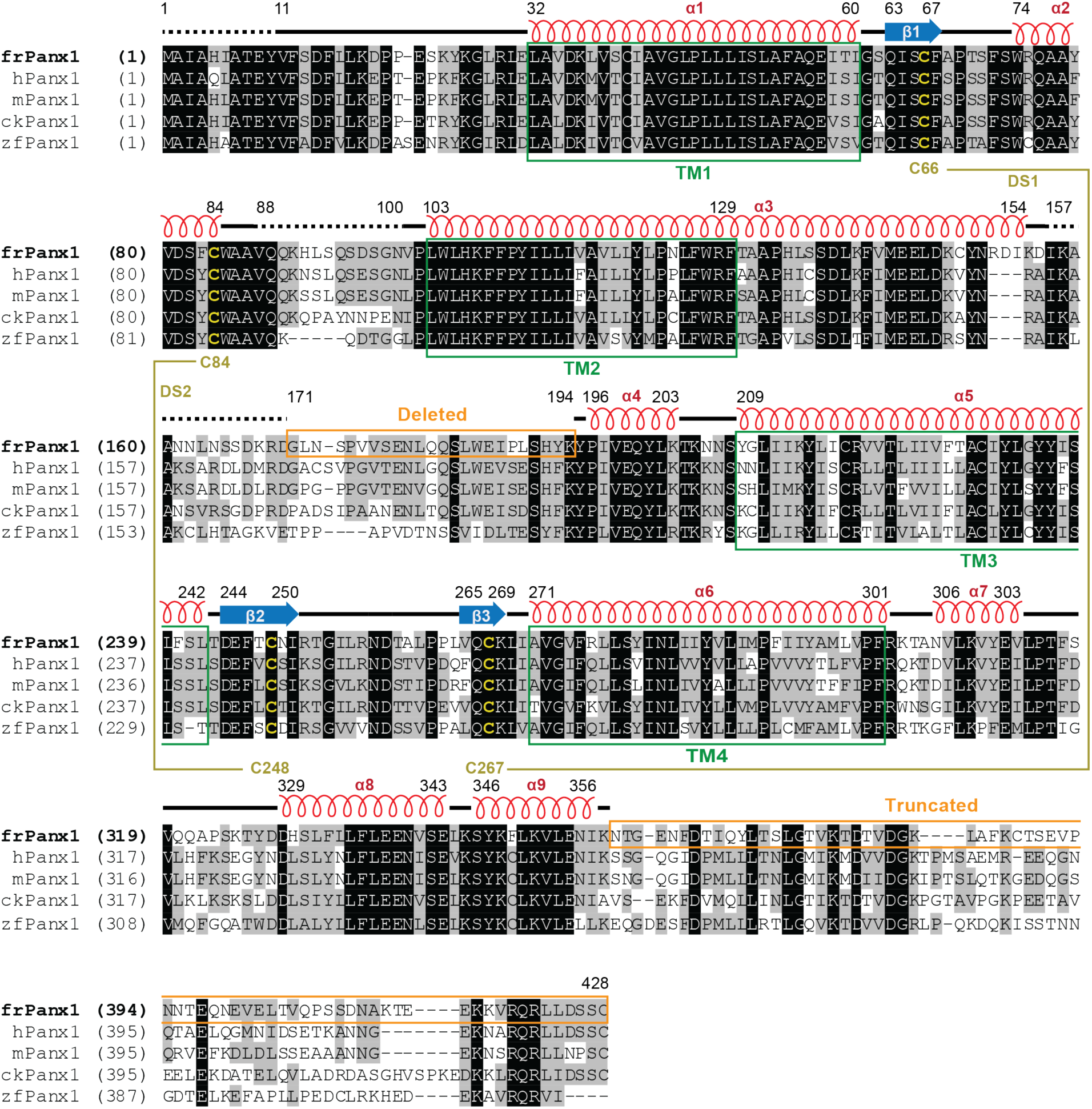
Sequence alignment and structural features. Amino acid sequence of frPanx1 compared to various Panx1 orthologues. Amino acids highlighted in black are completely conserved, in grey are similar, and in white are not conserved. Disulfide-forming cysteines are highlighted in yellow. Secondary structure features are shown above the sequence alignment. Green boxes depict the transmembrane helix boundaries, and orange boxes show deleted or truncated regions to create frPanx1-ΔLC.

**Figure1-figure supplement 2.**
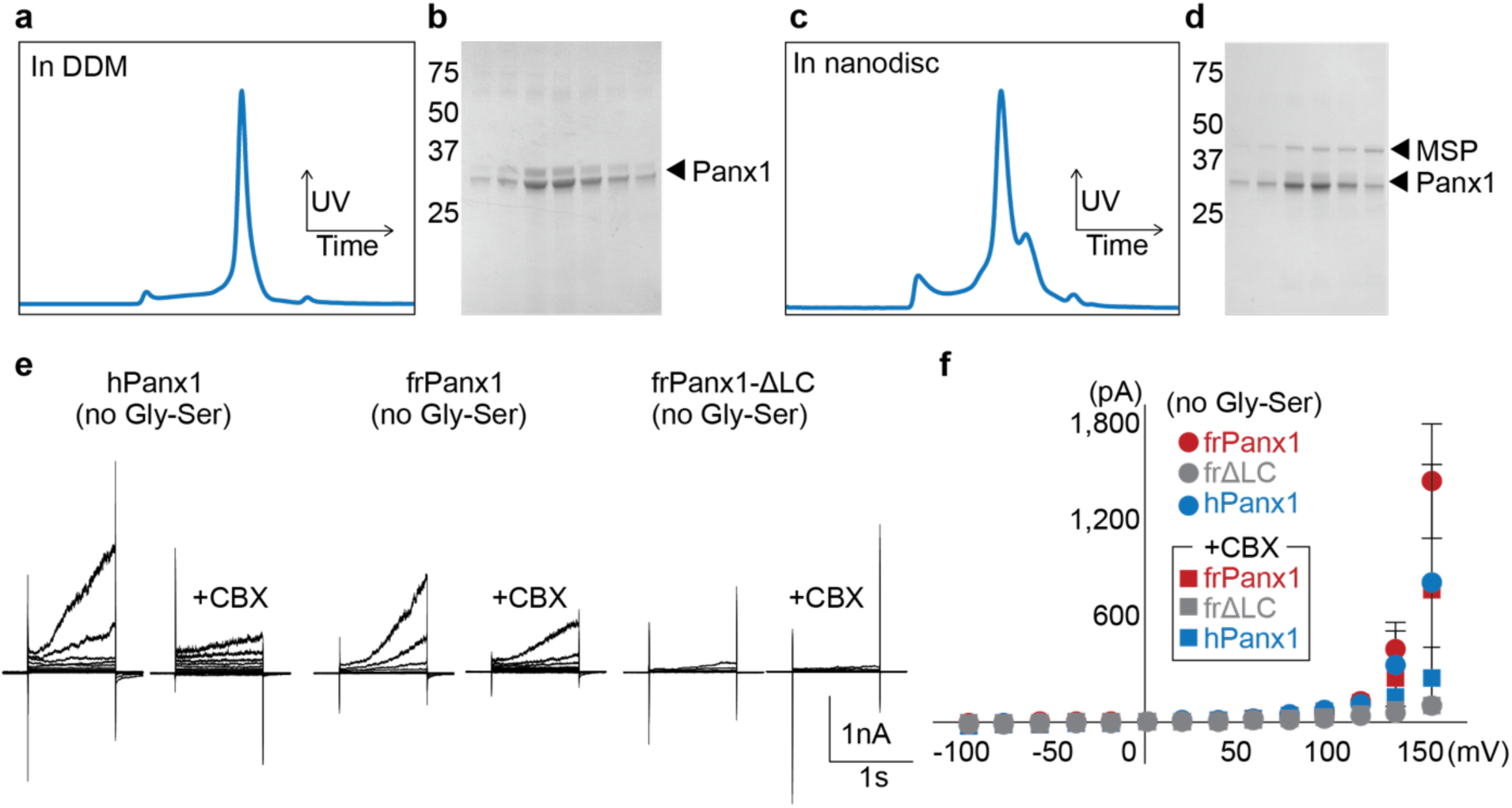
Characterization of frPanx1-ΔLC. **a**, Size exclusion chromatogram of frPanx1-ΔLC. Concentrated protein was injected onto a Superose 6 10/300 column equilibrated with 150 mM NaCl, 10 mM Tris pH 8.0, 1 mM EDTA, 0.5 mM DDM. **b**, SDS-PAGE analysis of peak fractions collected from **a. c**, Size exclusion chromatogram of frPanx1-ΔLC after reconstitution into nanodiscs. The running buffer contained 150 mM NaCl, 10 mM Tris pH 8.0, 1 mM EDTA. **d**, SDS-PAGE analysis of peak fractions collected from **c. e**, Whole-cell recordings of wild-type (no Gly-Ser) hPanx1, frPanx1, and frPanx1-ΔLC. Whole-cell patches from transfected HEK 293T cells were obtained, held at −60 mV, and stepped between −100 mV and +160 mV for 1 s. CBX (100 μM) was applied through a rapid solution exchanger. **f**, Current-voltage plot of wild-type pannexin recordings shown in **e**. Each circle represents the mean current at a particular voltage, with squares depicting the same current when treated with 100 μM CBX. N=3-11.

**Figure1-figure supplement 3.**
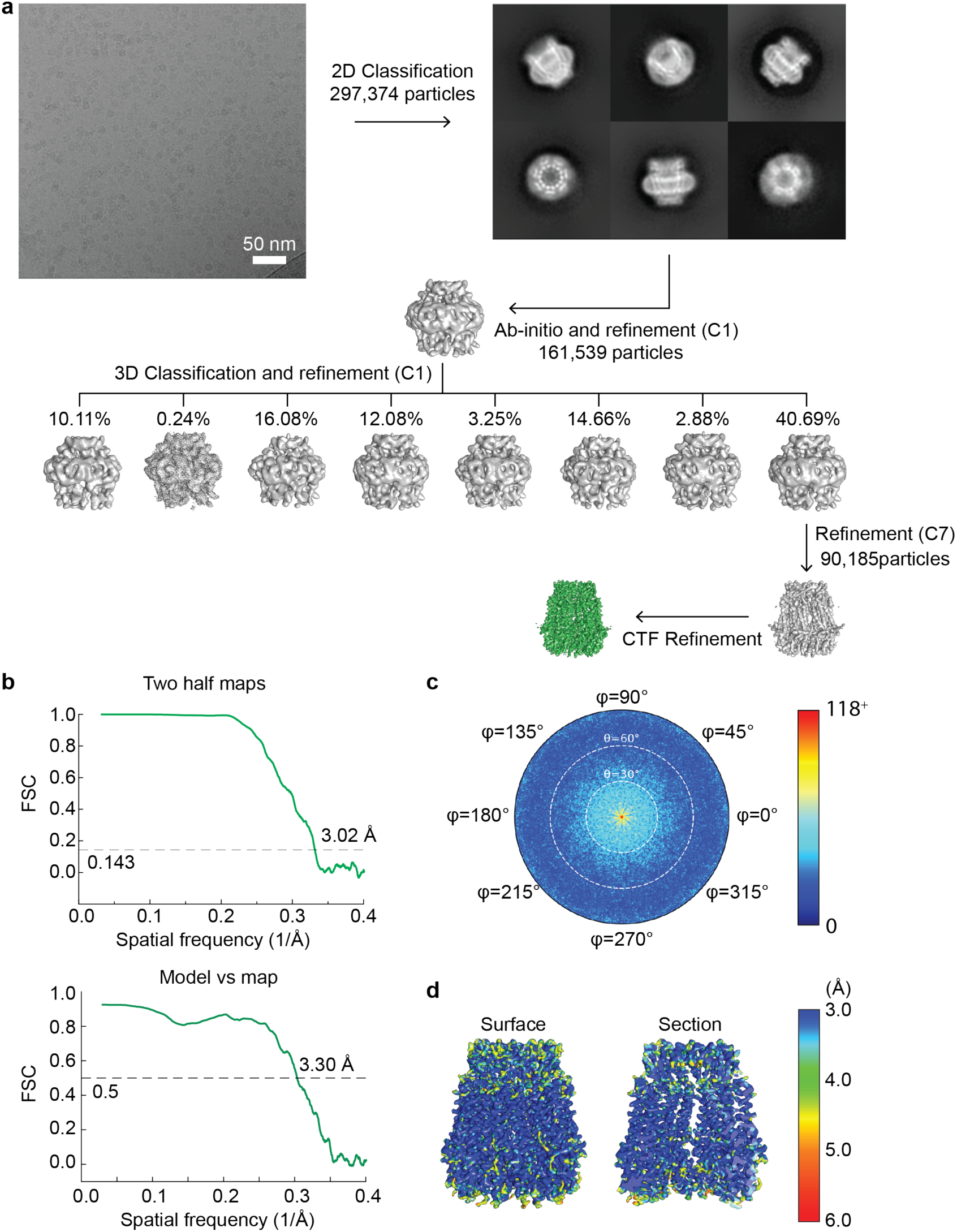
Cryo-EM image processing workflow for single particle analysis of frPanx1-ΔLC. **a.** A representative micrograph (scale bar = 50 nm), representative 2D class averages, and the 3D classification workflow are shown. **b.** The FSC plots of the two half maps (top) and the map vs model (bottom) are shown. **c.** The angular distribution plot for class 3. **d.** Local resolutions of class 3 were calculated using ResMap (Kucukelbir et al., 2014).

**Figure1-figure supplement 4.**
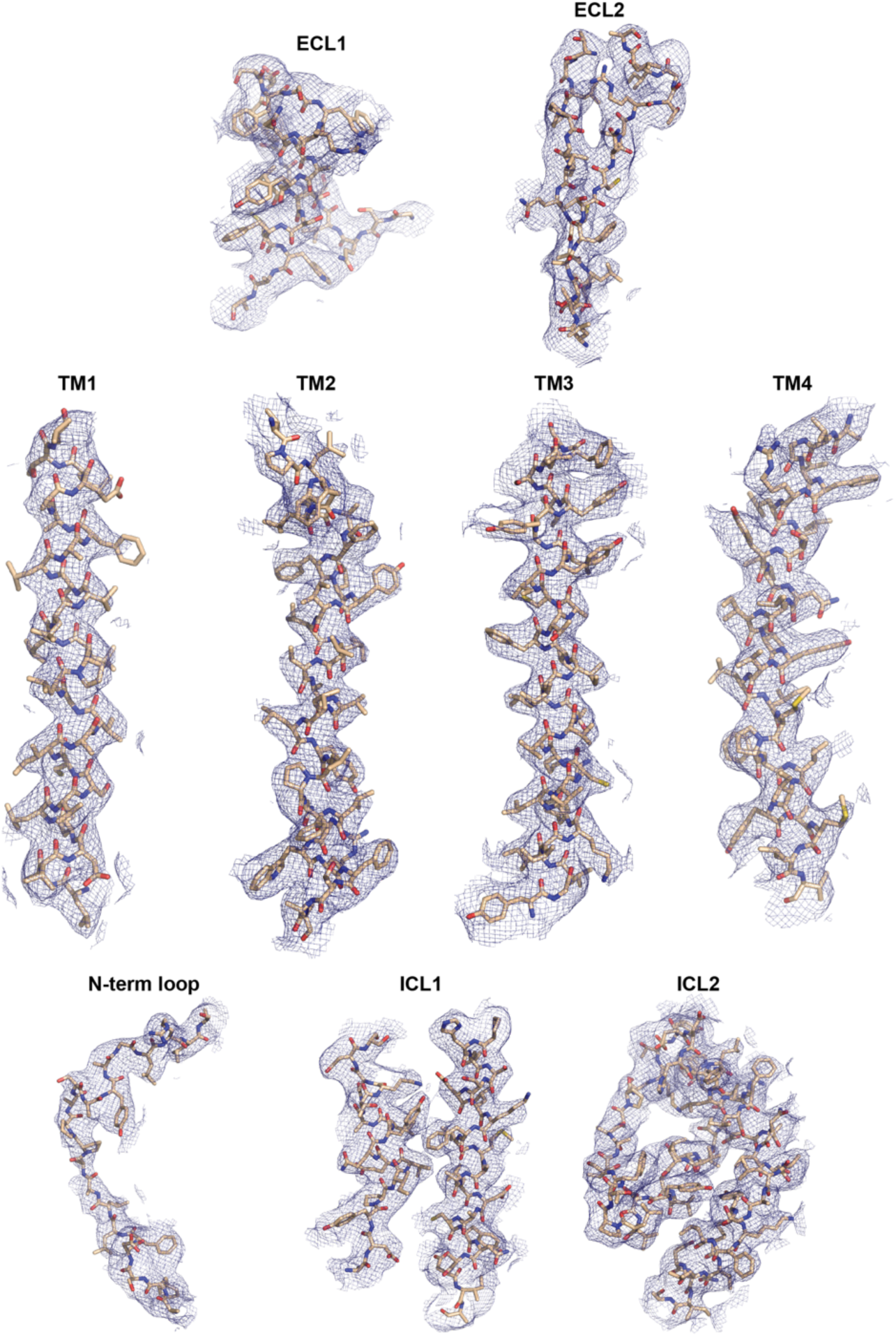
Representative cryo-EM density of frPanx1-ΔLC. Each domain is shown as stick representation and fit into the corresponding density contoured at σ=4.5.

**Figure2-figure supplement 1.**
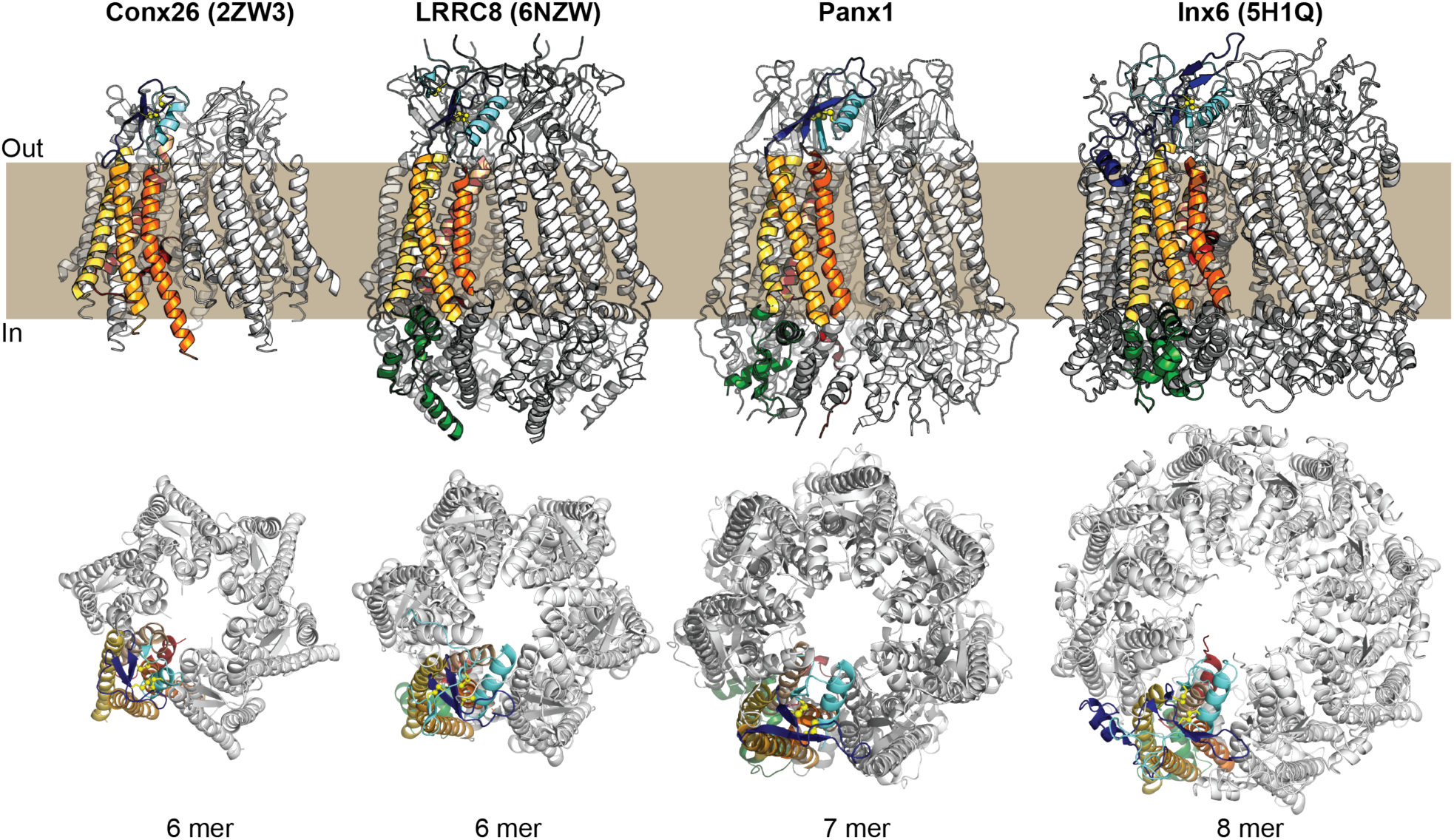
Comparison of frPanx1 with other large pore channels. The structures of connexin-26 (PDB: 2ZW3), LRRC8 (PDB: 6NZW), Panx1, and innexin-6 (PDB: 5H1Q) are shown from within the plane of the membrane (top) and viewed from the extracellular side (bottom). One subunit of each channel is shown colored with transmembrane domains in orange/yellow, extracellular loops in blue, and intracellular domains in green as shown in Fig. 2a.

**Figure2-figure supplement 2.**
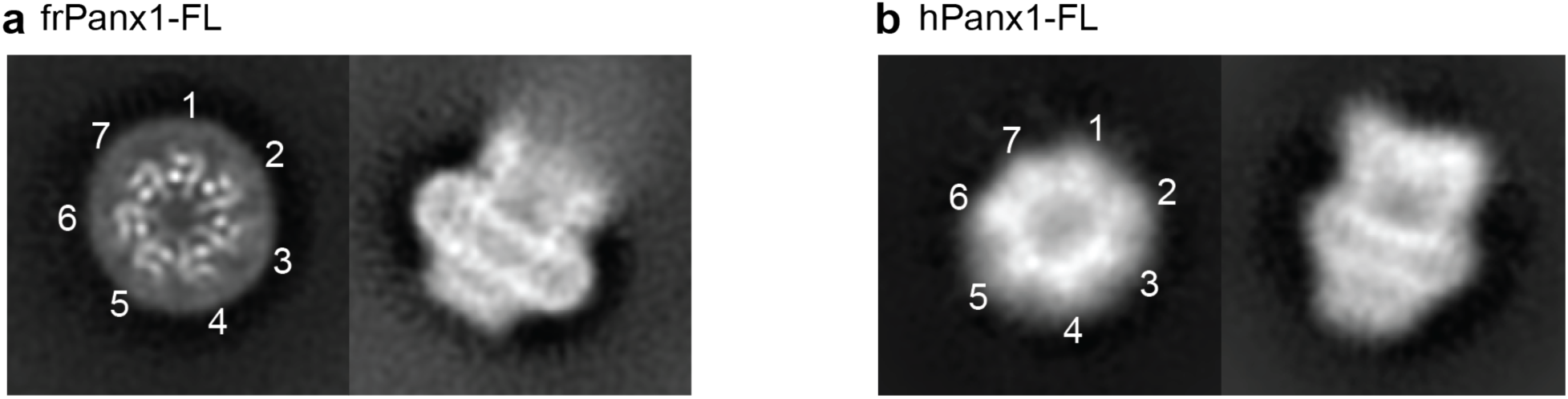
2D classes of full-length frog and human pannexin 1. **a**. 2D classes of the full-length frog pannexin1 in nanodiscs showing top (left) and side (right) views. All seven protomers are labeled with numbers in the 2D class of the top view. **b.** 2D classes of the full-length human pannexin 1 in DDM in the similar orientation to *panel a*. The top view shows a heptameric assembly (numbered). No symmetry was imposed in 2D classification.

**Figure2-figure supplement 3.**
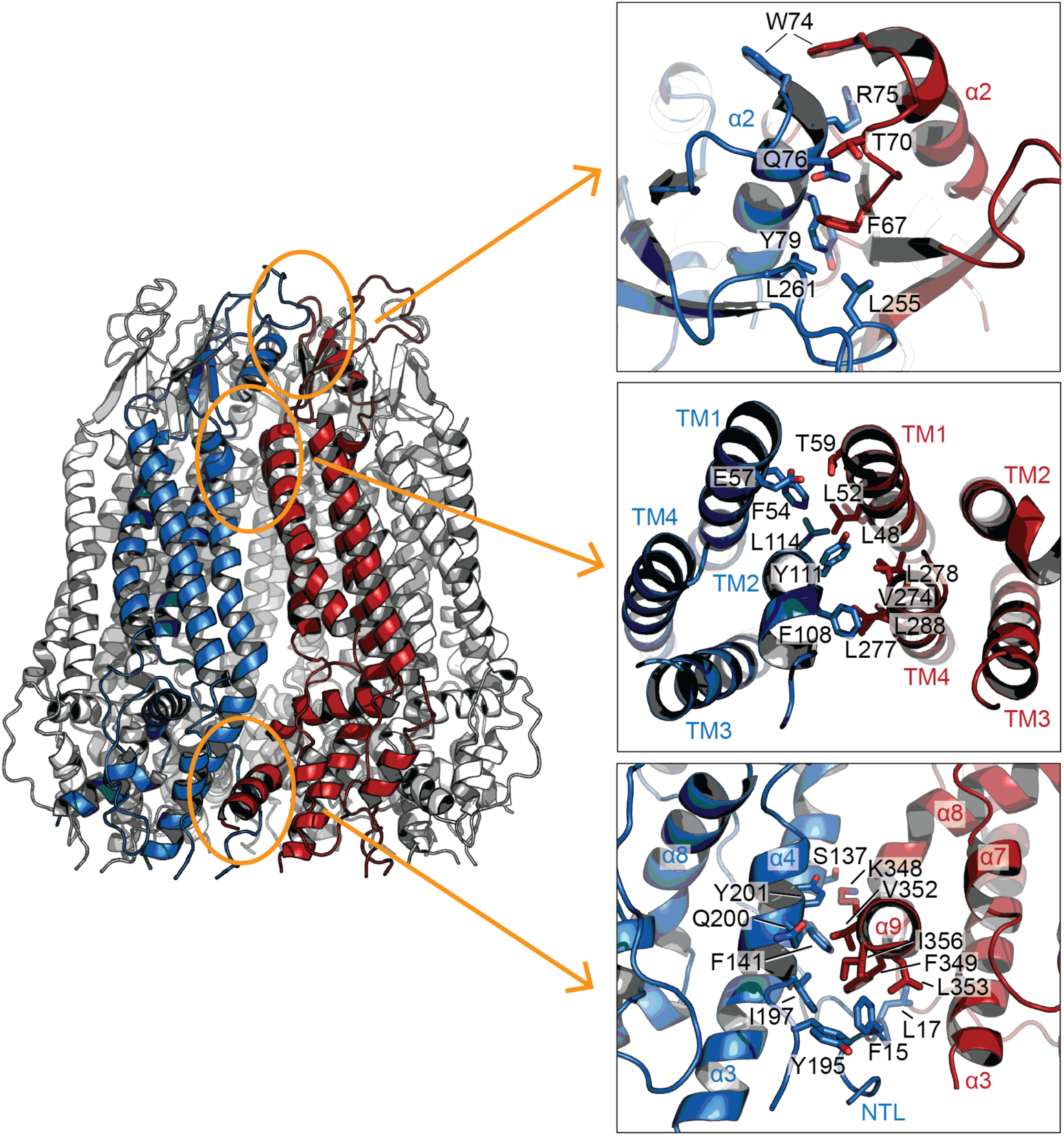
Inter-subunit interactions. Three major inter-subunit interfaces between two neighboring subunits (blue and red) are highlighted in orange ovals (left). Close-up views (right) show the highly-conserved residues mediating the inter-subunit interactions.

**Figure3-figure supplement 1.**
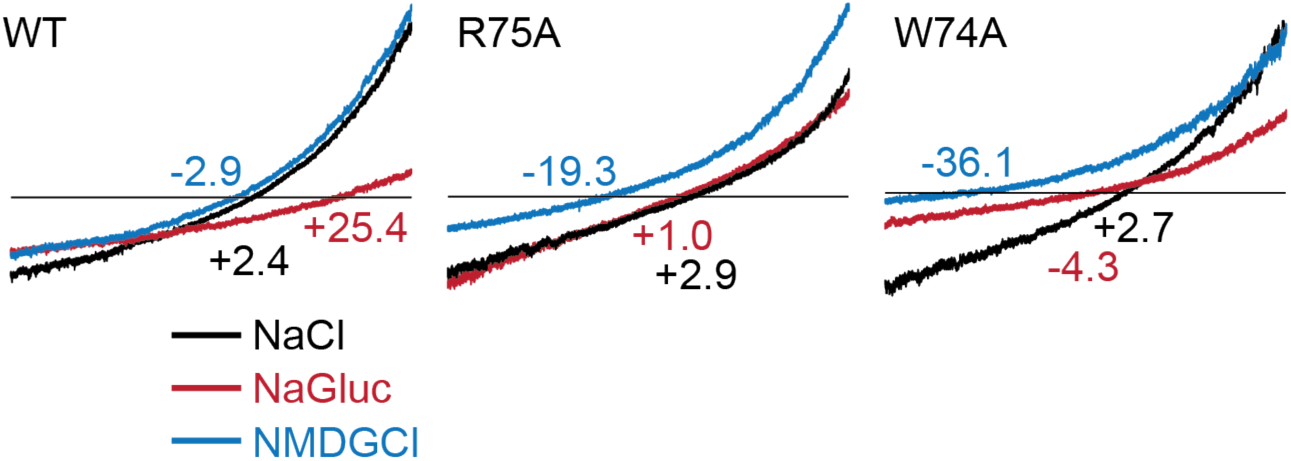
Representative traces of the ramp recordings. HEK293 cells were held at −60 mV and ramped between −100 mV and + 100 mV over 3s duration. The numbers indicate the reversal potentials in mV.

## Methods

### Purification of frPanx1-ΔLC

frPanx1 (NP_001123728.1) was synthesized (Genscript) and cloned into the BamHI/ XhoI sites of pCNG-FB7 vector containing a C-terminal Strep-tag II (WSHPQFEK). Amino acids from the IL1 and IL2 were removed by standard PCR strategies, and the BamHI site was also removed by quickchange mutagenesis. The full length frPanx1 and hPanx1 (NP_056183.2; synthesized by Genscript) were also subcloned into pCNG-FB7 vectors by standard PCR. Sf9 cells were infected with high titer baculovirus (20-25 mL P2 virus/ L cells) at a cell density of 2.5-3.0×10^6^ cells/ mL and cultured at 27 °C for 48 hours. Cells were collected by centrifugation, washed once with PBS, and lysed by nitrogen cavitation (4635 cell disruption vessel; Parr Instruments) at 600 psi in PBS containing leupeptin (0.5 μg/mL), aprotinin (2 μg/mL), pepstatin A (0.5 μg/mL), and phenylmethylsulfonyl fluoride (0.5 mM). Broken cells were centrifuged at 12,000 x g for 10 minutes, and membranes were collected by ultracentrifugation at 185,000 x g for 40 minutes. Membranes were suspended and solubilized in PBS containing 1% C12E8 (Anatrace) for 40 minutes, followed by ultracentrifugation at 185,000 x g for 40 minutes. Solubilized material was incubated with StrepTactin Sepharose High Performance resin (GE Healthcare) for 40 minutes in batch. Resin was collected onto a gravity column (Bio-Rad), washed with 10 column volumes of wash buffer (150 mM NaCl, 100 mM Tris-HCl pH 8.0, 1 mM EDTA, 0.5 mM C12E8), and eluted with 5 column volumes of wash buffer supplemented with 2.5 mM desthiobiotin. Eluted protein was concentrated and further purified on a Superose 6 10/300 Increase column (GE Healthcare) with 150 mM NaCl, 10 mM Tris pH 8.0, 0.5 mM DDM as the running buffer. Peak fractions were collected and pooled. All steps were performed at 4 °C or on ice.

### Reconstitution into nanodiscs

MSP2N2 apolipoprotein was expressed and purified as described previously (Ritchie et al., 2009), and the N-terminal His tag was cleaved off using TEV protease prior to use. To incorporate frPanx1 into nanodiscs, soybean polar extract, MSP2N2 and frPanx were mixed at final concentrations of 0.75, 0.3 and 0.3 mg/ml, respectively. The mixture was incubated end-over-end for 1 hour at 4 °C, followed by detergent removal by SM2 Bio-Beads (Bio-Rad). The supernatant and wash fractions were collected after an overnight incubation (∼12 hours) and further purified by size exclusion chromatography using a Superose 6 10/300 column in 20 mM Tris-HCl pH 8.0, 150 mM NaCl, 1 mM EDTA. Peak fractions were pooled and concentrated to 3 mg/mL.

### Cryo-EM sample preparation and image collection

frPanx1 in nanodiscs or hPanx1 in n-Dodecyl-β-D-Maltopyranoside (DDM; Anatrace) were applied to glow-discharged lacey carbon coated copper grids (Electron Microscopy Services). The grids were blotted for 4 s with blot force 7 at 85% humidity at 15 °C, and plunge frozen into liquid ethane using a Vitrobot Mark IV (Thermo Fisher). All data were collected on a FEI Titan Krios (Thermo Fisher) operated at an acceleration voltage of 300 keV. For frPanx1-ΔLC, a total of 2034 images were collected at 130k magnification with a pixel size of 1.07 Å in electron counting mode. Each micrograph was composed of 32 frames collected over 4 s at a dose of 1.79 e / Å^2^ / frame and a total exposure per micrograph of 57.3 e / Å^2^. Data were collected using EPU software (FEI). For full-length frPanx1 in nanodiscs, a total of 574 images were collected at 130k magnification with a pixel size of 1.06 Å in electron counting mode. Each micrograph was composed of 50 frames collected over 10 s at a dose of 1.4 e / Å^2^ / frame. The total exposure per micrograph was 70 e / Å^2^. Data were collected using SerialEM (Schorb et al., 2019). Data for full-length hPanx1 in DDM were collected in a similar fashion.

### Cryo-EM image processing and single particle analysis

Warp was used for aligning movies, estimating the CTF and particle picking for frPanx1-ΔLC and full-length hPanx1. For full-length frPanx1, movie alignment and CTF estimation were performed using the program Unblur and CTFFind, respectively, within the cisTEM package (Grant et al., 2018). 2D classification, ab-initio 3D map generation, 3D refinement, 3D classification, per particle CTF refinement and B-factor sharpening were performed using the program cisTEM (Grant et al., 2018). The single particle analysis workflow for frPanx1-ΔLC is shown in Extended Data Fig. 3. De novo modeling was performed manually in Coot (Emsley and Cowtan, 2004). The final model was refined against the cryo-EM map using PHENIX real space refinement with secondary structure and Ramachandran restraints (Adams et al., 2010). The FSCs were calculated by phenix.mtriage. Data collection and refinement statistics are summarized in Extended data Table 1.

### Electrophysiology

HEK293 cells were plated onto 12-mm glass coverslips (VWR) in wells of a 6-well plate and transfected 24 hours later with 500-800 ng plasmid DNA using FUGENE 6 (Promega) according to the manufacturer’s instructions. Recordings were performed ∼16-24 hours later using borosilicate glass micropipettes (Harvard Apparatus) pulled and polished to a final resistance of 2-5 MΩ. Pipettes were backfilled with (in mM) 147 NaCl, 10 EGTA, 10 HEPES pH 7.0 with NaOH. Patches were obtained in external buffer composed of (in mM) 147 NaCl, 10 HEPES pH 7.3 with NaOH, 13 glucose, 2 KCl, 2 CaCl_2_, 1 MgCl_2_. A rapid solution exchange system (RSC-200; Bio-Logic) was used to perfuse cells with CBX or various salt solutions. Currents were recorded using an Axopatch 200B amplifier (Axon Instruments), filtered at 2 kHz (Frequency Devices), digitized with a Digidata 1440A (Axon Instruments) with a sampling frequency of 10 kHz, and analyzed with the pClamp 10.5 software (Axon Instruments). For voltage step recordings, Panx1 expressing cells were held at −60 mV and stepped to various voltage potentials for 1 s in 20 mV increments before returning to −60 mV. For ramp recordings, cells were held at −60 mV, and ramped between −100 mV and + 100 mV over 3s duration.

## Acknowledgements

We thank the members of the Kawate lab for discussions. This work was supported by the National Institutes of Health (GM114379 to T.K.; NS113632 to H.F.; GM008267 to K.M., E.K., and J.K; GM008267 to K.M.), Robertson funds at Cold Spring Harbor Laboratory, Doug Fox Alzheimer’s fund, Austin’s purpose, and Heartfelt Wing Alzheimer’s fund (to H.F.). J.L.S. is supported by the Charles H. Revson Senior Fellowship in Biomedical Science.

## Author information

**Kevin Michalski**

Present address: WM Keck Structural Biology Laboratory, Cold Spring Harbor Laboratory, Cold Spring Harbor, NY, USA

**Julia Kumpf**

Present address: Brookline High School, 115 Greenough Street, Brookline, MA, USA

## Contributions

K.M. generated expression constructs, purified proteins, and optimized protein samples for cryo-EM. J.S. prepared the nanodisc samples for cryo-EM, collected cryo-EM data, and proceeded for 3D reconstruction. E.H. performed electrophysiology experiments. J.K. identified and optimized frPanx1 expression constructs. H.F. collected and processed cryo-EM data, analysed the data, and wrote the manuscript. T.K. designed the experiments, analysed the data, and wrote the manuscript.

